# RegKnock: identifying gene knockout strategies for microbial strain optimization based on regulatory and metabolic integrated network

**DOI:** 10.1101/438168

**Authors:** Zixiang Xu

**Affiliations:** National Engineering Laboratory for Industrial Enzymes and Tianjin Engineering Center for Biocatalytic Technology, Tianjin Institute of Industrial Biotechnology, Chinese Academy of Sciences, Tianjin 300308, China; Key laboratory of systems microbial biotechnology, Tianjin Institute of Industrial Biotechnology, Chinese Academy of Sciences, Tianjin 300308, China

**Keywords:** metabolic network, regulatory network, integrated network, mixed integer bi-level linear programming (MIBLP), strain optimization, flux balance analysis, constraint-based modeling, gene knockout, gene deletion, metabolic engineering, systems biology, synthetic biology

## Abstract

**Background:** Gene knockout has been used to improve the conversion ratio of strains for some chemical products. Based on mixed integer bi-level linear programming (MIBLP) and cell network models, there have been several algorithms to predict the target for deletion to improve the productivity of chemicals. At present, the cell models on which these algorithms based have changed from metabolic network to metabolic-regulatory integrated network, for integrated network is more comprehensive in describing the behavior of cells. Metabolic-regulatory integrated network is better than metabolic network in flux prediction, but will introduce integer variables in the inner of MIBLP. How to solve the intractable MIBLP, however, is not explicated clearly as in mathematical literatures, especially for MIBLP with integer variables in the inner problem (named as MIBLP-2) where integer variables are introduced by the flux balance analysis (FBA) for integrated network. Dual theory was still be used to transform MIBLP-2 to a single level with ignoring integer variables in the inner problem. Intelligent computation is another choice for solving MIBLP, but it usually was used to solve the single level nonlinear programming (NLP) which was the transformation from MIBLP by using joint objective of upper/lower level, while the equivalence between this MIBLP and this NLP was not be proved in mathematics.

**Methods:** In this study, we develop a new target predicting algorithm for gene knockouts, named RegKnock. The cell model on which we base is metabolic-regulatory integrated network as well. When solving the MIBLP-2, RegKnock uses Parallel Genetic Algorithm (PGA), but not use joint objective. GA was used to generate control variables of the upper, indicating which genes should be deleted, while the fitness function is to maximize the objective product calculated from the inner FBA of the integrated network. FBA of the inner problem, a mixed integer programming, could be solved by existing optimization softwares. Parallel computation aims to accelerate finding the optimal solution and thus decreases the time of computation.

**Results and Conclusions:** With comparing with OptORF and OptFlux, two published target predicting algorithm for gene knockouts which also aiming at integrated network, two merits have been shown for RegKnock, i.e. absolutely accuracy and not a long time of computation. So RegKnock is a nice algorithm for predicting algorithm for gene deletions as for integrated network.

## Introduction

The concepts and methods of metabolic engineering have been elucidated in the 1990s [Gregory, 1991; J. E. Bailey, 2001]. In this century, metabolic engineering has been pinned outstanding hopes on energy, environment and climate improvement, food sources and human health, and so on [Betenbaugh, 2008]. The technique of DNA recombinant makes it possible to manipulate genetic changes, and gene knockout method has been used to improve the conversion ratio of industrial strains for some chemical products. With the development of system biology and synthetic biology and especially with constraint-based modeling [Price ND, 2003] and flux balance analysis (FBA) [Blazeck, 2010], by utilizing genome-scale cell network models, metabolic engineering has come into the era of system metabolic engineering [Orth JD, 2010].

There are a series of published algorithms to predict the targets for deletion [Burgard, 2003; Pharkya, 2004; Kiran, 2005; Pharkya, 2006; Miguel, 2008; Lun DS, 2009; Tepper, 2010; Sridhar, 2010; Kim, 2010; Rocha, 2010; Paulo, 2011; Xu, 2013]. Bi-level optimization, which was introduced first time by OptKnock [Burgard, 2003], is the core of these algorithms. Bi-level optimization can solve the conflict of cell growth and maximum bioengineering objective. For the control variables in these algorithms indicated which genes should be knockout, the upper-level variables are integer or binary, so these bi-level optimization models belong to the type of MIBLP (mixed integer bi-level linear programming) that is difficult in solving [Paulo, 2011]. There are three keys for these predicting methods for gene deletion targets. 1) The first is which kind of cell model, metabolic network or integrated network (metabolic/regulatory), is based on? Of course, metabolic and regulatory integrated network is more comprehensive than metabolic network. Up to now, genome-scale integrated network can refer to rFBA [Markus, 2003 & 2002] and SR-FBA [Shlomi, 2007]. If using metabolic network, the inner problem of MIBLP is a FBA model and thus is a linear programming (LP), and we can name it as the type of MIBLP-1; If using integrated network, the inner problem of MIBLP is rFBA or SR-FBA model and thus will include integer variables to describe the states of genes and proteins, so is a mixed integer programming (MIP) as itself, and we can name this kind of MIBLP as the type of MIBLP-2. 2) The second is how to solve MIBLP? Two main methods have been applied, i.e. numerical and intelligent computation [Zeynep, 2005]. Numerical computation here means transforming the MIBLP to a single level and then solves it with existing optimization softwares, while intelligent computation here means using Evolutionary Algorithms (EAs), Simulated Annealing (SA), Genetic Algorithms (GA), etc. 3) The third is about the performance of predicting algorithms, such as solution time (long-L/moderate-M/short-S), solution number (just one or multi if existed), and knockout number (fixed number or a searching scope), etc. **Table 1** has given a statistic on those published algorithms [Burgard, 2003; Pharkya, 2004; Kiran, 2005; Pharkya, 2006; Miguel, 2008; Lun DS, 2009; Tepper, 2010; Sridhar, 2010; Kim, 2010; Rocha, 2010; Paulo, 2011; Xu, 2013] which predicting the targets for deletion.

**Table 1.** statistic on those published algorithms to predict deletion targets

| Algorithm Name | Cell Model | Mathematic Model | Solving Method | Deletion Number | Solution Number | Solution Time | Published Time |
| --- | --- | --- | --- | --- | --- | --- | --- |
| OptKnock | Metabolic network | MIBLP-1 | Transform to MILP<br>Solved by optimization softwares | fixed | one | Moderate | [Burgard, 2003] |
| OptStrain | Metabolic network | MIBLP-1 | Transform to MILP<br>Solved by optimization softwares | fixed | one | not tried | [Pharkya, 2004] |
| OptGene | Metabolic network | MIBLP-1 | Using EA to generate control variables Fitness function is only cell growth | fixed | one | Long | [Kiran, 2005] |
| OptReg | Metabolic network | MIBLP-1 | Transform to MILP<br>Solved by optimization softwares | fixed | one | not tried | [Pharkya, 2006] |
| meta-heuristics | Metabolic network | MIBLP-1 | Transform to NLP with joint objective<br>Solved by EA/SA | scope | one | not tried | [Miguel, 2008] |
| GDLS | Metabolic network | MIBLP-1 | Transform to MILP<br>Solved by optimization softwares | fixed | one | Moderate | [Lun DS, 2009] |
| RobustKnock | Metabolic network | MIBLP-1 | Transform to MILP<br>Solved by optimization softwares | fixed | one | Long | [Tepper, 2010] |
| ReacKnock | Metabolic network | MIBLP-1 | Transform to MILP<br>Solved by optimization softwares | scope | multi | Moderate | [Xu, 2013] |
| OptForce | Metabolic network | MIBLP-1 | Transform to MILP<br>Solved by optimization softwares | fixed | one | not tried | [Sridhar, 2010] |
| OptORF | Integrated network | MIBLP-2 | Transform to MILP, the inner is not SR-FBA<br>Solved by optimization softwares | fixed | one | not tried | [Kim, 2010] |
| OptFlux | Metabolic network<br>Integrated network | MIBLP-1<br>MIBLP-2 | Transform to NLP with joint objective<br>Solved by EA/SA | scope | multi | Long | [Rocha, 2010;<br>Paulo, 2011] |
| RegKnock | Integrated network | MIBLP-2 | Solve MIBLP directly, the inner is SR-FBA<br>Solved by Parallel Genetic Algorithm (PGA) | scope | multi | Moderate | This study |

About model base for predicting method of gene deletion targets, from the perspective of development, metabolic and regulatory integrated network is better than metabolic network, because integrated network is more comprehensive in describing the behavior of cells [Markus, 2003 & 2002; Shlomi, 2007]. At the same time, with the difference of applied model, the targets predicted are different as well. If it is for metabolic network, the targets predicted are reactions or enzymes [Burgard, 2203]; if it is for integrated network, the targets predicted are genes [Kim, 2010; Rocha, 2010]. As for the complex gene-protein-reaction mapping, gene engineering favors gene targets much more [Kim, 2010; Rocha, 2010]. About solving MIBLP, it seems all these published algorithms to predict the deletion targets seldom refer not to mathematical literatures that are strict in deduction and verification for how to solving a MIBLP. OptKnock [Burgard, 2003] was the first and also the base of many other algorithms (RoboustKnock, GDLS, OptORF and so on), which regarded the control variables of the upper problem as parameters, transformed the inner problem to its dual form, and finally got a single level one, a mixed integer linear programming (MILP). Although it was an advance for OptORF to utilize integrated network model [Kim, 2010], the inner problem for the MIBLP-2 of this method should be SR-FBA and should include integer variables to describe the states of genes and proteins, but its inner problem actually was FBA and it did not explicit clearly how to cope with those integer variables involving in the states of genes and proteins. The integers in the inner problem should not be moved to the outer, so it is difficult to solve MIBLP-2 [Scott, 2011]. OptFlux that also based on integrated network transformed MIBLP-2 to a single level nonlinear programming (NLP) by using joint objective, Biomass-Product Coupled Yield (BPCY) or Product Yield with Minimum Biomass (PYMB) [Rocha, 2010], and then utilized EAs or SA to solve the NLP [Rocha, 2010]. But whether the equivalence between its MIBLP-2 and its NLP was not be proved in mathematics. OptFlux can also be applied to metabolic network models [Rocha, 2010], where the targets predicted are reactions. OptGene aimed to metabolic network models and also use Evolutionary Algorithm to solve the MIBLP-1 [Kiran, 2005]. But unlike OptFlux, EA in OptGene is just used to generate control variables of the upper and its inner problem was FBA while the fitness function was cell growth [Kiran, 2005]. A shortcoming of OptGene was a long time of computation.

In this study, we develop a new predicting algorithm of gene knockout targets. The cell model on which we base is metabolic and regulatory integrated network (i.e. SR-FBA) and the mathematic frame thus is MIBLP-2, illustrating in **Fig 1** and named as RegKnock. When solving this MIBLP-2, RegKnock uses Genetic Algorithm (GA). We adopt the strategy from OptGene rather than from OptFlux, i.e. we use GA to generate control variables of the upper rather than transform MIBLP-2 to a single level nonlinear programming (NLP) by using joint objective. The inner problem of our MIBLP-2 is still SR-FBA, while the fitness function is to maximize the objective product calculated from the inner FBA of the integrated network. We develop the GA algorithm with Matlab rather than use its standard GA toolbox. In order to accelerate shrink speed, we introduce Parallel Genetic Algorithm (named as PGA) and this saves the computational time greatly. In every generation of GA of RegKnock, we save the better solution of a given number, so we can obtain multi solutions at last. Multi solutions have the advantage in the choice of gene operation. As for the gene knockout algorithms to integrated network, we have done some case studies comparing with OptFlux and OptORF. We have mentioned the shortcomings of OptFlux and OptORF above. As for RegKnock, first of all, it absolutely obeys the inner SR-FBA model (integrated network), that is to say, the predicted result of RegKnock is correct for inner SR-FBA model. If we removed those target genes from the inner SR-FBA model, the results of cell growth and industrial objective will be equal to the predicted values of the lower level objective and the upper-level objective. Then if SR-FBA model is correct, the predicted result of RegKnock is correct. The comparing results from RegKnock with OptORF and OptFlux have supported RegKnock in its accuracy (according to with inner SR-FBA) and nice computational speed of PGA.

**Fig 1.**
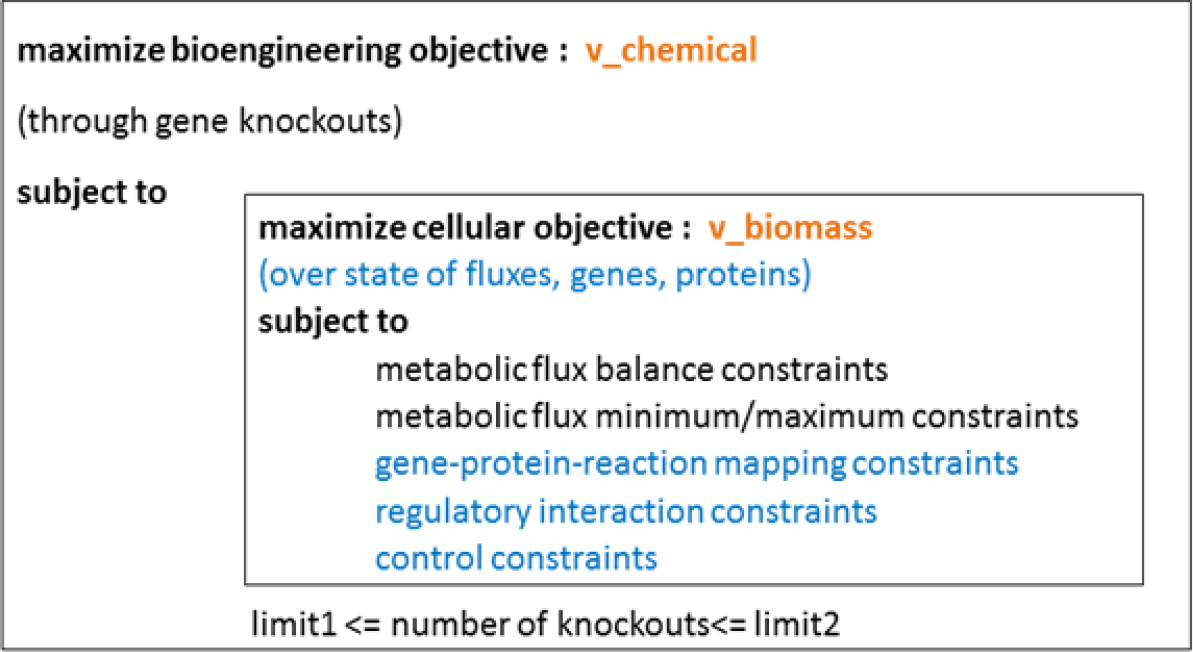
The MIBLP frame of RegKnock: the cell model is based on integrated network, targets are genes which are also control variables, the knockout number is a scope between two given numbers, and the inner problem is a SR-FBA. Both inner and outer problem have the integer variables, so it is the MIBLP-2 type, most difficult to solve.

## Methods

### Metabolic/regulatory integrated network

rFBA was the first frame work to integrate the metabolic and regulatory network in cells [Markus, 2003 & 2002], where Boolean logic was used to describe regulatory network. Based on rFBA model, Shlomi et al [Shlomi, 2007] have provided a MILP frame work, named as SR-FBA which can predict the behavior of the cell, including cell growth, gene expression states, protein expression states, and fluxes through reactions. Its mathematic model is as follows

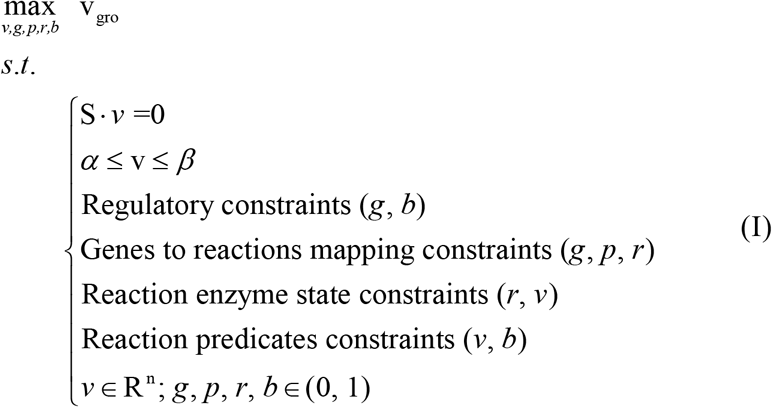

*v* representing fluxes through reactions, *g* representing the Boolean expression state of all genes, *p* representing the presence of each protein, *r* representing the presence of a catalyzing enzyme for each reaction, *b* representing the state of each flux predicate, *S* is the stoichiometric matrix, *α* and *β* represent lower and upper bounds on the flux reaction rates.

### Mathematical model of RegKnock

The main idea of Regknock is also a bi-level optimization in MIBLP-2 type. The upper level is for industrial objective and the inner problem is SR-FBA. So the deletion targets that Regknock predict are genes rather than reactions/enzymes, and the cell model on which Regknock base is metabolic and regulatory integrated network in SR-FBA frame [Shlomi, 2007].

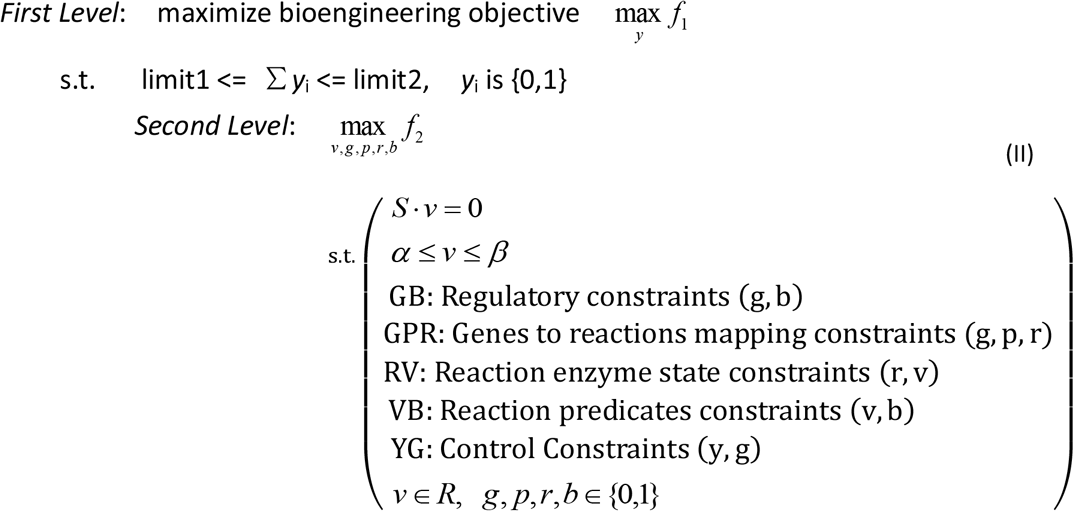

Here, *f*_1_ is the objective function to maximize industrial production, *f*_2_ is the objective function for cells to maximize the growth, i.e. *v*gro; *S* · *v* = 0 is the kinetic equations, and are minimum and maximum of flux in every reaction; *v*, *g*, *p*,*r*, *b* respectively are flux, gene state, protein state, reaction state, flux predicate (decision of a flux being larger than a value); *y* is the control variable, y(i)=0 means the gene should be deleted. We can rewrite the above model strictly in mathematical format.

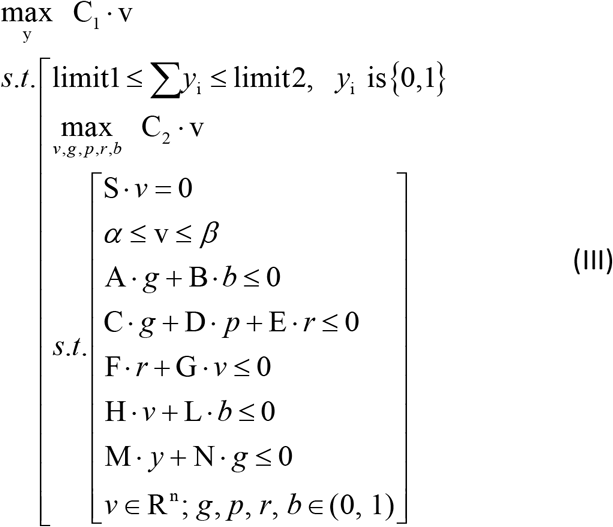

Here, all the variable vectors have been stated above; A ^~^ L, C_1_, C_2_ are matrixes in proper dimensions.

### Method to solve the MIBLP-2 model

In order to solve the above mathematic MIBLP-2 model (II) or (III), we didn’t transform this MIBLP-2 to a single level nonlinear programming (NLP) by using joint objective. Actually, we used GA to generate control variables of the upper, while the inner problem (SR-FBA, a MILP) could be solved by commercial optimization software, such as Gurobi [http://www.gurobi.com]. The objective of the inner problem of the GA is cell growth, but the fitness function of the GA is to maximize the objective product. Here we didn’t use standard GA toolbox of Matlab, otherwise, we developed a new GA algorithm with Matlab intending to solve MILP especially. We introduce parallel computation to accelerate shrinking speed, so the core of our solving method is PGA (Parallel Genetic Algorithm). At the same time, as for a MIBLP, its solution may be multiply. Multi solutions have the advantage in the choice of gene operation as well. In order to obtain multi solutions of the MIBLP-2 (II)/ (III), in every generation of PGA of RegKnock, we save the better solution of a given number. But for OptFlux, it gets the multi solutions through multi runs, i.e. obtaining one solution in one time of computation. As illustrated in **Fig 2**, we give the flow chart of the PGA for RegKnock.

**Fig 2.**
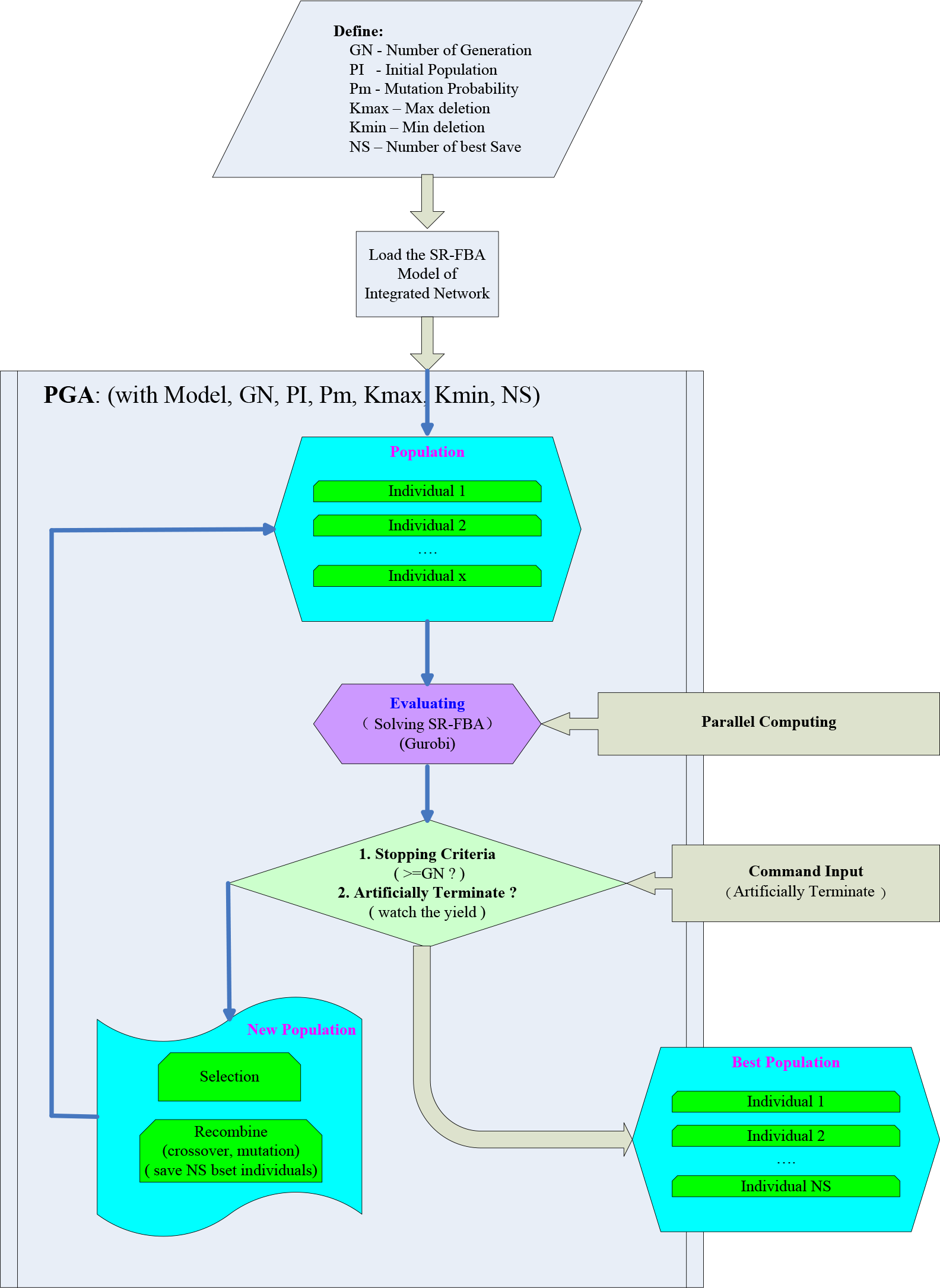
Flow chart of the PGA for RegKnock

### Verification method

Whenever we solve a MIBLP with numerical methods or intelligent methods, we should verify its solution. The best way is to substitute the solution, the values of control variables from the upper level that indicate which genes should be deleted, to the inner problem and calculate the inner problem again. Compare the value of cell growth after removing these target genes with the value of an objective function of the first level that we predicted previously; Compare the flux rate of produced objective chemical after removing these target genes with the value of an objective function of the lower level that we predicted previously. If they are equal respectively, the solution is correct.

## Results and Discussion

### Case study 1: Comparison between RegKnock and OptORF / OptFlux for producing Ethanol with *E.coli*

We used the same metabolic and regulatory integrated network of *E. coli_*iJR904 [Reed JL, 2003; Covert, 2004] which was also been used in OptORF and OptFlux as case studies. Shlomi et al. have given its SR-FBA model [Shlomi, 2007] that was used here in our computation as the presentation of integrated network. 5-gene knockouts for Ethanol production have been predicted by OptORF [Kim, 2010] and OptFlux [Paulo, 2011] and we also gave the prediction of 5 gene knockouts for Ethanol with RegKnock. The substrate was glucose, its input rate was 10 mmol/gDW/hr and minimal media was in anaerobic conditions. Growth was set to no less than 0.1 mmol/gDW/hr. We firstly testified the predicted strategies of OptORF and OptFlux, i.e. we removed those predicted genes from the SR-FBA model (setting their corresponding gene variables to 0) and did FBA again. The predicted strategies and predicted flux values of OptORF and OptFlux were respectively from the references [Kim, 2010; Paulo, 2011]. OptORF and OptFlux have not done FBA testification in their case studies. We found that the output rates of Ethanol from both OptORF and OptFlux were actually much smaller than the predicted values in every deletion strategy, as shown in **Table 2**. We then used RegKnock to predict 5-gene knockouts for Ethanol production with *E. coli* and RegKnock return 5 strategies for 5-gene knockouts. The first two got to the almost theoretical value of Ethanol production, and especially the predicted values of growth and the chemical producing rate had no error in FBA testification, as shown in **Table 2**. At the same time, the fluxes of Ethanol production predicted by OptORF and OptFlux were far from the theoretical value. Growth rate and chemical producing rate of wildtype *E. coli* and the maximum yield have been provided in **Table 2** as well.

**Table 2.** Testification of OptORF / OptFlux on the 5 gene knockouts and comparison with RegKnock: the original literature data of growth and chemical producing rate were under the 18.5 input rate of glucose [Kim, 2010, Paulo, 2011], and here we translated them into the input rate of glucose in 10 mmol/gDW/hr.

| Chemical<br>(condition) | Algorithm | Knockouts (5 genes with the<br>names) | Biomass<br>rate | Chemical<br>rate | FBA testification<br>(Biomass / Chemical) |
| --- | --- | --- | --- | --- | --- |
| Ethanol<br>(glc= -10,<br>anaerobic) | Wild_Type |  | 0.2146 | 8.3244 |  |
|  | Max_Yield |  |  | 20.000 |  |
|  | OptORF | fnr gntR pflB tdcE pgi | 0.1216 | 17.24 | 0, 10 |
|  |  | arcA gntR pta eutD tpiA | 0.1319 | 17.86 | 0, 8.9995 |
|  |  | fnr gntR pflB tdcE tpiA | 0.127 | 18.1 | 0, 10 |
|  | OptFlux | gnd atpF purU purT arcA | 0.236 | 14.5589 | 0.2146, 8.3244 |
|  |  | atpG speF purU arcA zwf | 0.236 | 14.5589 | 0.2146, 8.3244 |
|  |  | atpI purU arcA zwf | 0.236 | 14.5589 | 0, 10 |
|  |  | focA fruK arcA focB | 0.2251 | 16.6494 | 0.2146, 8.3244 |
|  |  | atpE purU arcA zwf | 0.236 | 14.5589 | 0, 10 |
|  |  | yqeA atpE purU arcA zwf | 0.236 | 14.5589 | 0.2146, 8.3244 |
|  |  | pgi fabH atpG thrB proA | 0.0542 | 16.8293 | 0.2146, 8.3244 |
|  |  | pgi fabH atpG mtlR thrB | 0.0543 | 16.828 | 0.2146, 8.3244 |
|  |  | ptsI gdhA gnd folD atpB | 0.1736 | 17.2472 | 0.2146, 8.3244 |
|  |  | ptsI gdhA gnd folD atpA | 0.1736 | 17.2472 | 0.2146, 8.3244 |
|  |  | ptsI gdhA atpI zwf deoC | 0.1785 | 17.2343 | 0, 10 |
|  |  | ptsH gdhA folD atpE zwf | 0.1736 | 17.2472 | 0.2146, 8.3244 |
|  |  | ptsH gdhA folD atpG zwf | 0.1736 | 17.2472 | 0, 10 |
|  |  | focA atpI pntB deoD focB | 0.0686 | 18.5169 | 0.2035, 12.5005 |
|  |  | pgi fabH atpI thrB proA | 0.0542 | 16.8293 | 0.2146, 8.3244 |
|  |  | ptsI gdhA folD atpG zwf | 0.1736 | 17.2472 | 0, 10 |
|  |  | ptsI gdhA gnd folD atpA | 0.1736 | 17.2472 | 0.2146, 8.3244 |
|  |  | ptsH gdhA folD atpA zwf | 0.1736 | 17.2472 | 0.2146, 8.3244 |
|  | RegKnock | lytB sfuA gnd menB malT atpG | 0.1984 | 14.2885 | 0.1984, 14.2885 |
|  |  | gnd menB malT atpG deoC | 0.1984 | 14.2885 | 0.1984, 14.2885 |
|  |  | caiF gnd menB malT atpG deoC | 0.1984 | 14.2885 | 0.1984, 14.2885 |
|  |  | yeaV atpE wecE thiG aceA | 0.2025 | 12.9085 | 0.2025, 12.9085 |
|  |  | PdxA citD cmk gapC_2 aceA | 0.2134 | 8.3340 | 0.2134, 8.3340 |

### Case study 2: Target prediction of gene knockout strategies for producing different chemicals with *E.coli*

We utilized RegKnock to predict different chemicals produced by *E. coli*, including Succinate, Formate, D-Lactate, and Threonine. The integrated network model was still SR-FBA model of *E. coli* [Shlomi, 2007]. The minimal media was also glucose and its input rate was 10 mmol/gDW/hr. D-Lactate and Threonine productions were in aerobic conditions while others were in anaerobic condition. In each case, RegKnock would return one strategie for about 12-gene knockouts. All the predicted knockout strategies and their corresponding FBA testification were shown in **Table 3**. As a comparison, **Table 3** has also given with growth rate and chemical producing rate of wild type *E. coli* under the same growth condition and the maximum yield that *E. coli* can produce. From **Table 3**, we can see that in most cases, the output rates of produced chemicals predicted by RegKnock have got to a higher yield than the wild-type to the same chemicals. Especially and above all, all the predicted knockouts absolutely accorded with their corresponding FBA testification. If we accredited that the SR-FBA model could rightly describe the behavior of *E. coli*, the predicted results of RegKnock were correct.

**Table 3.** Target predictions of about 12 gene knockouts for producing different chemicals

| Chemical (condition) | Algorithm | Knockouts (5 genes with the b numbers) | Biomass rate | Chemical rate | FBA testification (Biomass rate / Chemical rate) |
| --- | --- | --- | --- | --- | --- |
| Succinate (glc= -10, anaerobic) | Wild_Type |  | 0.2146 | 0.0789 |  |
|  | Max_Yield |  |  | 11.5363 |  |
|  | RegKnock | b0080 b0619 b0620 b1602 b1605 b2029<br>b2411 b2465 b3366 b3386 b3930 b3938 | 0.0016 | 3.9907 | 0.0016, 3.9907 |
| Formate (glc= -10, anaerobic) | Wild_Type |  | 0.2146 | 17.5627 |  |
|  | Max_Yield |  |  | 24.2954 |  |
|  | RegKnock | b0080 b0348 b1441 b1521 b1761 b1852<br>b2283 b3566 b3748 b3907 b4125 | 0.0016 | 19.9816 | 0.0016, 19.9816 |
| D-Lactate (glc= -10, aerobic) | Wild_Type |  | 0.2146 | 0 |  |
|  | Max_Yield |  |  | 19.9997 |  |
|  | RegKnock | b0171 b0662 b0809 b0828 b0972 b1380<br>b1702 b1761 b2239 b2781 b3112 b3973 | 0.1873 | 6.3699 | 0.1873, 6.3699 |
| Threonine (glc= -10, aerobic) | Wild_Type |  | 0.2146 | 0 |  |
|  | Max_Yield |  |  | 12.3361 |  |
|  | RegKnock | b0080 b0350 b0401 b0729 b1474 b1761<br>b1852 b2787 b2920 b2979 b3111 b3803<br>b4229 | 0.0016 | 1.9838 | 0.0016, 1.9838 |

The optimal production did not converge to near theoretical yield while it has reached to the final generation. The parameters of GA, such as the number of population, the number of generations and mutation rate, will affect the optimal production predicted. So if we hope to obtain a better optimal solution, we should set a larger number of generations and population. For the reason of computational time, in this case, we just show the effectiveness of our algorithms but do not want to obtain the practical strategies of strain optimization, so we just set a small number of generations and population, respectively to be 200 and 500. Another aspect for affecting the optimal production is the number of gene knockouts. A very small number of gene knockouts will not make it easy to get a high yield of objective chemicals. The following case will show the relationship between the predicted yield and the number of gene knockouts.

### Case study 3: The relationship between the predicted yield and the number of gene knockouts

We utilized RegKnock to predict the product of Ethanol produced by *E. coli*. The computation condition is the same as in case 2. In this case, RegKnock would return the strategies for the number of 2 to 6 in gene knockouts and we listed out those better strategies. All the predicted knockout strategies and their corresponding yields were shown in **Table 4**. As we said above, a very small number of gene knockouts will not make it easy to get a high yield of objective chemicals. With the number of gene knockouts increasing, the predicted yield will increase under the same computation condition (hardware and software, parameter setting, minimal media setting).

**Table 4.** The relationship between the predicted yield and the number of gene knockouts

| Chemical (condition) | Algorithm | Knockouts (3 to 6 genes with the b numbers) | Biomass rate | Chemical rate |
| --- | --- | --- | --- | --- |
| Ethanol (glc= -10, anaerobic) | Wild_Type |  | 0.2146 | 8.3244 |
|  | Max_Yield |  |  | 20.000 |
|  | RegKnock | b0007 b0099 b0125 b0243 b0469 b4025 | 0.0522 | 15.9153 |
|  |  | b0062 b0099 b0125 b0243 b0469 b4025 | 0.0522 | 15.9153 |
|  |  | b0061 b0099 b0125 b0243 b0469 b4025 | 0.0522 | 15.9153 |
|  |  | b0099 b0125 b0243 b0469 b4025 | 0.0522 | 15.9153 |
|  |  | b0052 b0678 b4025 b4040 | 0.0524 | 15.9138 |
|  |  | b2094 b2988 b3997 b4025 | 0.0523 | 15.9137 |
|  |  | b0485 b3726 b3731 | 0.2025 | 12.9058 |
|  |  | b0062 b1898 b3738 | 0.2025 | 12.9058 |
|  |  | b0080 b0173 b2805 | 0.2136 | 8.6977 |
|  |  | b0080 b2028 | 0.2136 | 8.6977 |

### Parameter setting of GA, computational time and algorithm convergence

In above case studies of RegKnock, we set the number of generations and the number of population respectively to be 200 and 500, set the mutation probability to be 0.45, and run the studies on a server with 48 cores.

In general, the computational time of GA will take a longer time than numerical methods. The computational time of each case in this study was less than 24 hours. Of course, the number of population and the number of generations will affect the solving time, the larger the number, the longer the computational time, and the higher the yields we predicted. At the same time, parallel computing has greatly decreased the time of computation. The solver of optimization will also affect the computational time and Gurobi was utilized in our study.

Algorithm convergence was good for case study 1 - 3. In all the cases, the optimal production reached to a higher value with the increasing of the number of generations. **Fig 3** showed the value change of optimal production over the number of generations for case study 3 on gene knockouts for D-Lactate.

**Fig 3.**
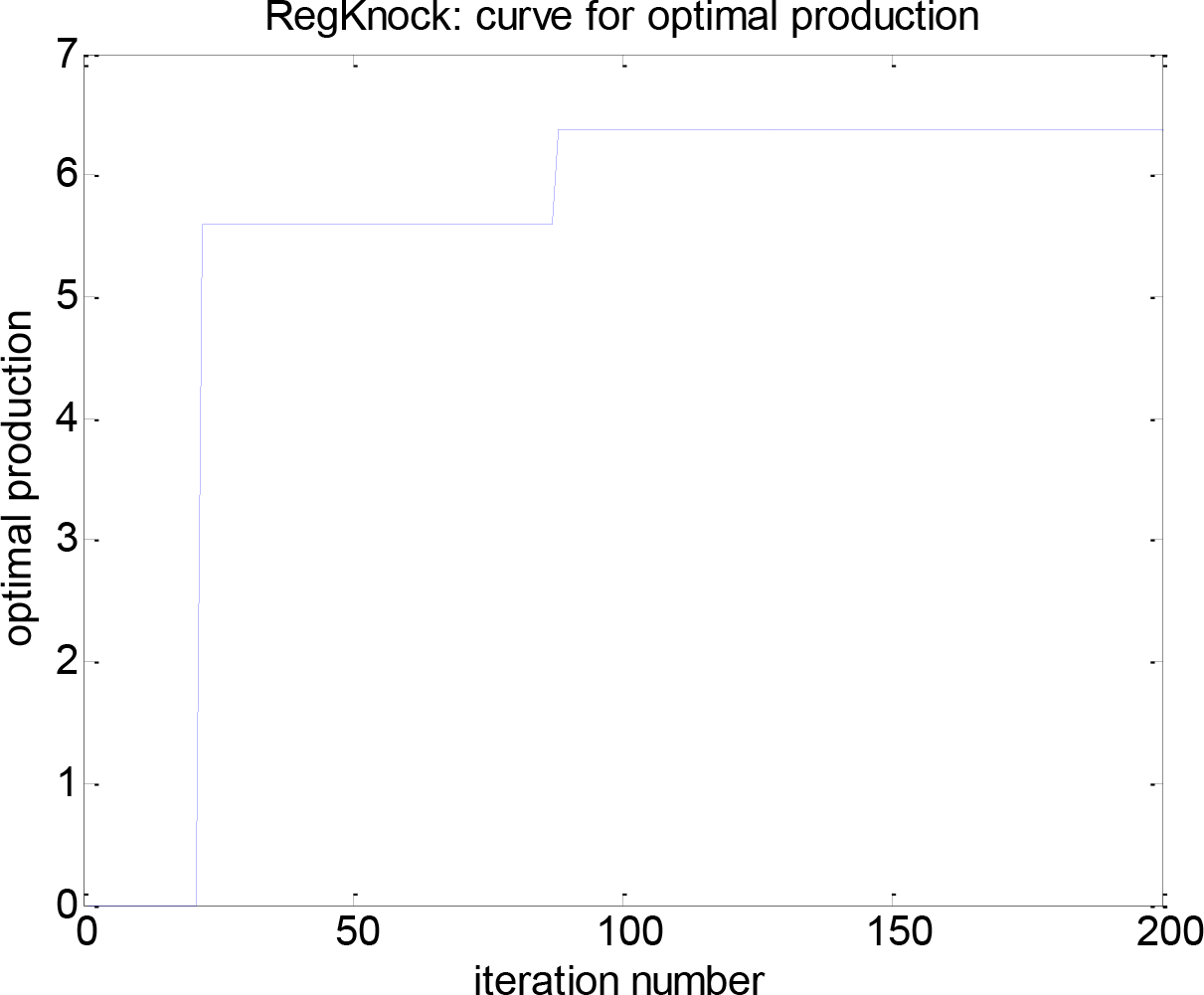
The value change of optimal production over the number of generation for case study 1 by using RegKnock to producing D-Lactate with *E.coli*. x-axis was the number of generation and y-axis was the flux producing D-Lactate.

## Conclusions

DNA recombinant is the main tool of optimizing industrial strains for some chemical products. In order to reduce blindness of gene knockouts to improve the conversion ratio, we should utilize those tools of computational biology to predict deletion targets. For metabolic-regulatory integrated networks are more comprehensive in describing the behavior of cells, we should base our predicting algorithms on this kind of cell models. Several previous algorithms of this type have been designed alike, but they were not proved in mathematics for solving MIBLP-2 which is the frame work of the predicting methods aiming to integrated network. FBA testification has shown their unreliability.

In this study, we design an algorithm for predicting deletion targets to improve the conversion ratio, named RegKnock, which is also based on the cell models of integrated networks. RegKnock uses Genetic Algorithm to solve MIBLP-2 and at the same time, it introduces parallel computation to speed up the solving procedure. RegKnock has performed several merits over those previous alike algorithms: 1) It is absolutely reliable and its predicted deletion targets absolutely accord with FBA testification, for it utilizes GA directly; 2) It need relatively less time to find suboptimal or optimal solution, for it introduces parallel computation; 3) It could return multiple solutions thus increase the choices in DNA recombinant experiments.

## Availability

The Matlab code of RegKnock algorithm could be provided by email requirement.

## Acknowledgements

Support for this work was provided by “National Natural Science Foundation of China (31370829, 31370113)’’, “Tianjin Research Program of Application Foundation and Advanced Technology (15JCYBJC23600)”, “Tianjin Science and Technology Committee (11ZCZDSY08600)”.

## Author contributions

Conceived and designed the experiments: ZX. Performed the experiments: ZX. Analyzed the data: ZX QW DZ JS. Contributed reagents/materials/analysis tools: QW DZ JS. Wrote the paper: ZX.

